# IRG1 and iNOS act redundantly with other interferon gamma-induced factors to restrict intracellular replication of *Legionella pneumophila*

**DOI:** 10.1101/731182

**Authors:** Jordan V. Price, Daniel Russo, Daisy X. Ji, Roberto Chavez, Lucian DiPeso, Angus Yiu-Fai Lee, Jörn Coers, Russell E. Vance

## Abstract

Interferon gamma (IFNγ) restricts the intracellular replication of many pathogens, but how IFNγ confers cell-intrinsic pathogen resistance remains unclear. For example, intracellular replication of the bacterial pathogen *Legionella pneumophila* in macrophages is potently curtailed by IFNγ, but consistent with prior results, no individual genetic deficiency we tested compromised IFNγ-mediated control. Intriguingly, however, we observed that the glycolysis inhibitor 2-deoxyglucose (2DG) partially rescued *L. pneumophila* replication in IFNγ-treated macrophages. 2DG inhibits glycolysis and triggers the unfolded protein response, but unexpectedly, it appears these effects are not responsible for perturbing the antimicrobial activity of IFNγ. Instead, we found that 2DG rescues bacterial replication predominantly by inhibiting the induction of two key antimicrobial factors, inducible nitric oxide synthase (iNOS) and immune responsive gene 1 (IRG1). Using immortalized and primary macrophages deficient in iNOS and IRG1, we confirm that loss of both iNOS and IRG1, but not individual deficiency in each gene, partially reduces IFNγ-mediated restriction of *L. pneumophila*. Further, using a combinatorial CRISPR/Cas9 mutagenesis approach, we find that mutation of iNOS and IRG1 in combination with four other genes (CASP11, IRGM1, IRGM3 and NOX2) results in a total loss of *L. pneumophila* restriction by IFNγ in primary bone marrow macrophages. There are few, if any, other examples in which the complete set of cell-intrinsic factors required for IFNγ-mediated restriction of an intracellular bacterial pathogen have been genetically identified. Our results highlight the combinatorial strategy used by hosts to block the exploitation of macrophages by pathogens.

**Importance:** *Legionella pneumophila* is one example among many species of pathogenic bacteria that replicate within mammalian macrophages during infection. The immune signaling factor interferon gamma (IFNγ) blocks *L. pneumophila* replication in macrophages and is an essential component of the immune response to *L. pneumophila* and other intracellular pathogens. However, to date, no study has determined the exact molecular factors induced by IFNγ that are required for its activity. We generated macrophages lacking different combinations of IFNγ-induced genes in an attempt to find a genetic background in which there is a complete loss of IFNγ-mediated restriction of *L. pneumophila*. We successfully identified six genes that comprise the totality of the IFNγ-dependent restriction of *L. pneumophila* replication in macrophages. Our results clarify the molecular basis underlying the potent effects of IFNγ and highlight how redundancy downstream of IFNγ is key to prevent exploitation of the macrophage niche by pathogens.

## Introduction

Macrophages are preferred host cells for many species of intracellular bacterial pathogen. *Bona fide* pathogens of mammals, such as *Mycobacterium tuberculosis* (*Mtb*), *Listeria monocytogenes*, and *Salmonella enterica*, as well as environmental microorganisms that are “accidental” pathogens of mammals, such as *L. pneumophila*, display the ability to replicate efficiently in macrophages, demonstrating that these cells can provide a plastic niche suitable to the metabolic needs of distinct bacterial species (1). To defend against potential exploitation by diverse pathogens, including environmental microorganisms with which they have not co-evolved, macrophages require potent mechanisms to restrict intracellular bacterial replication. A cornerstone of the immune response to many intracellular pathogens is the cytokine interferon gamma (IFNγ). The importance of IFNγ is highlighted by the observation that genetic deficiencies in the IFNγ signaling pathway render humans highly susceptible to infections by intracellular pathogens, most notably *Mtb* and even normally benign environmental bacteria (2). Mice engineered to be deficient in the IFNγ pathway are also highly susceptible to intracellular bacterial pathogens, including *Mtb*, *L. monocytogenes*, *S. enterica*, *Brucella abortus*, and *L. pneumophila*, among others (3–9). Brown *et al* demonstrated that failure of IFNγ-deficient mice to control *L. pneumophila* likely occurs at the level of cell-intrinsic restriction of bacteria in monocyte-derived macrophages that infiltrate the lung following infection (10). Accordingly, *in vitro* infection models using bone marrow-derived macrophages (BMMs) have enabled meaningful study of the cell-intrinsic immune response to *L. pneumophila* coordinated by IFNγ. However, despite several decades of evidence supporting an essential role for IFNγ in the antimicrobial immune response, the precise mechanisms by which IFNγ acts to mediate cell-intrinsic control of *L. pneumophila* and other pathogens remain obscure.

Inducible nitric oxide synthase (iNOS, encoded by the gene *Nos2* in mice) plays a key role in the IFNγ-dependent response to *Mtb* and several other pathogens (11–13). iNOS facilitates the production of nitric oxide (NO), a toxic metabolite with direct antimicrobial activity. NO also acts as a regulator of host responses and coordinates metabolic changes in IFNγ-stimulated macrophages (14–16). While *Nos2*^−/−^ mice display increased susceptibility to infection by *Mtb*, it appears this is not simply due to direct cell-intrinsic antimicrobial effects of NO (17). In addition, the activity of iNOS is not absolutely required to control infection by many pathogens, suggesting that there are redundant iNOS-independent mechanisms that underlie the potency of IFNγ (18). Strikingly, while *L. pneumophila* does not display resistance to the effects of NO in broth, *Nos2*^−/−^ macrophages are not impaired in IFNγ-dependent restriction of *L. pneumophila* (19–21). This indicates either that *L. pneumophila* is resistant to the effects of iNOS/NO during infection or, more likely, that there are redundant factors induced by IFNγ that can restrict *L. pneumophila* in the absence of iNOS.

Previous work has attempted to address the possibility of redundancy in the IFNγ-dependent immune response to *L. pneumophila*. Pilla *et al* generated quadruple knockout (QKO) mice deficient in *Nos2*, *Cybb* (cytochrome b(558) subunit beta, encoding NADPH oxidase 2 aka NOX2), *Irgm1* (immunity-related GTPase family M member 1), and *Irgm3* (immunity-related GTPase family M member 3), all induced by IFNγ (20). NOX2 partners with phagosomal oxidase components to generate reactive oxygen species, which, like NO, can cause direct toxicity to phagocytized pathogens in neutrophils and macrophages (22, 23). IRGM1 and IRGM3 are antimicrobial GTPases that may participate in the disruption of membrane-bound, pathogen-containing compartments within phagocytes (20, 24). Remarkably, Pilla *et al* observed that macrophages derived from QKO mice retained potent restriction of *L. pneumophila* replication when stimulated with IFNγ, and further implicated the bacterial lipopolysaccharide detector caspase 11 (CASP11), which when activated can trigger host macrophage pyroptosis, in some of the residual IFNγ-dependent restriction of *L. pneumophila* replication in macrophages (20).

Recently, Naujoks *et al* implicated immune responsive gene 1 (IRG1, encoded by the gene *Acod1*) in the IFNγ-dependent immune response to *L. pneumophila*, demonstrating that driving *Acod1* expression in macrophages was sufficient to suppress *L. pneumophila* replication (21). However, the study did not address whether macrophages deficient in IRG1 were impaired in the ability to restrict *L. pneumophila* when stimulated with IFNγ. Like iNOS, IRG1 also generates a potentially toxic metabolite (itaconate), and contributes to metabolic changes that occur in inflamed macrophages (25–27).

*L. pneumophila* normally replicates in protozoan host amoebae, but can cause a severe pneumonia in humans, known as Legionnaires’ disease, through infection of lung macrophages. *L. pneumophila* employs a type-IV secretion system to translocate bacterial effector proteins into the host cytosol, allowing the bacteria to establish an intracellular replicative compartment (28). Flagellin produced by wild-type *L. pneumophila* can trigger host cell pyroptosis via the NAIP/NLRC4 inflammasome; however, *L. pneumophila* that lack flagellin (Δ*flaA*) are able to replicate to high levels in macrophages (29–35). We recently described a mutant strain of *L. pneumophila* (Δ*flaA*Δ*uhpC*) that is able to replicate in macrophages treated with 2-deoxyglucose (2DG), an inhibitor of mammalian glycolysis (36). This strain allows us to probe the role that host cell metabolism plays in the immune response to *L. pneumophila*.

In the present study, we use a combination of pre-existing knockout mouse models, pharmacological treatment with 2DG and other drugs, CRISPR/Cas9 genetic manipulation of immortalized mouse macrophages, and BMMs from novel strains of CRISPR/Cas9-engineered mice to survey of the factors required for IFNγ-dependent restriction of *L. pneumophila* in macrophages. Ultimately, we demonstrate that iNOS and IRG1 are sufficient and redundant in terms of IFNγ-dependent restriction of *L. pneumophila*. Further, we identify six IFNγ-inducible factors: iNOS, IRG1, CASP11, NOX2, IRGM1, and IRGM3, which are responsible for the entirety of the IFNγ-dependent restriction of *L. pneumophila* in macrophages.

## Results

### IFNγ restricts *L. pneumophila* replication across a spectrum of antimicrobial gene-deficient macrophages

In an attempt to identify specific factors that explain the ability of IFNγ to restrict *L. pneumophila* replication in macrophages to specific genetic factors associated with the immune response, we tested the ability of IFNγ to restrict *L. pneumophila* in BMMs derived from various knockout mice. Using an extensively validated strain of Δ*flaA L. pneumophila* that expresses luminescence (lux) genes from *P. luminescens* (20, 36–38), we confirmed that Δ*flaA L. pneumophila* replicates in unstimulated BMMs but does not replicate in BMMs stimulated with IFNγ (**Figure 1A**). As expected, BMMs lacking the IFNγ receptor (*Ifngr*^−/−^) do not restrict *L. pneumophila* replication in the presence of IFNγ (**Figure 1B**). We confirmed that BMMs lacking functional iNOS (*Nos2*^−/−^) retain IFNγ-dependent restriction of *L. pneumophila* (**Figure 1B**) (19–21). BMMs lacking MYD88 (*Myd88*^−/−^), a key adaptor in the innate inflammatory immune response triggered by bacterial pattern recognition, and BMMs lacking MYD88, NOD1, and NOD2 (*Myd88*^−/−^ *Nod1*^−/−^ *Nod2*^−/−^), which do not activate inflammatory NF-κB signaling in response to *L. pneumophila* (39), still restricted bacterial replication when stimulated with IFNγ (**Figure 1C**). Consistent with previous results (20), BMMs deficient in ATG5 (LysMCre^+^ *Atg5*^fl/fl^), a factor essential for autophagy (40), also mediate IFNγ-dependent restriction of *L. pneumophila* replication (**Figure 1D**). Pilla *et al* reported that guanylate binding proteins (GBPs) in conjunction with caspase-11 (CASP11) partially mediate IFNγ-mediated restriction of *L. pneumophila* in BMMs (20). We observed that BMMs from mice that lack a region of chromosome 3 containing five GBPs (GBP1, 2, 3, 5, and 7, *Gbp*^chr3−/−^) and also BMMs that that lack functional caspase-1 and CASP11 (*Casp1/11*^−/−^) largely retain the ability to restrict *L. pneumophila* in the presence of IFNγ (**Figure 1E**). Additionally, we observed that BMMs derived from mice lacking functional MYD88 and TRIF (*Myd88*^−/−^ *Trif*^−/−^), STING (*Goldenticket*) (41), IFNAR (*Ifnar*^−/−^), and TNF receptor (*Tnfr*^−/−^) all retained IFNγ-dependent restriction of *L. pneumophila* replication (data not shown). In sum, these data confirm that no single genetic factor studied to date is required to restrict *L. pneumophila* replication in macrophages stimulated with IFNγ.

**Figure 1.**
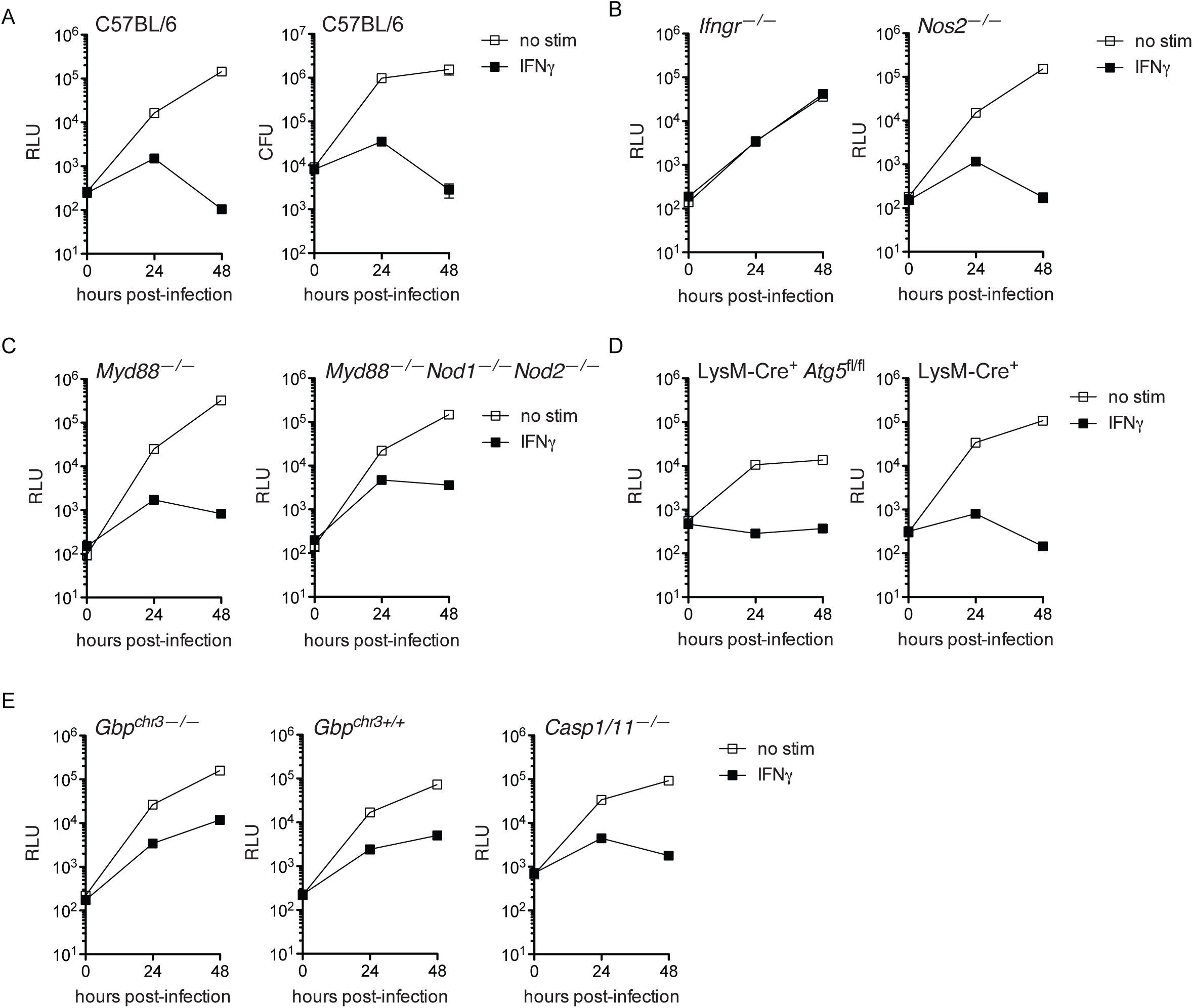
IFNγ restricts *L. pneumophila* replication across a spectrum of immune response-impaired macrophage genotypes. (**A**) Luminescence measured in relative light units (RLU, left) and recovery of colony forming units (CFU, right) of LP02 Δ*flaA* lux *pneumophila* from infected wild-type C57BL/6 BMMs either not stimulated (no stim) or stimulated with 6.0 ng/mL IFNγ. (**B**) RLU from LP02 Δ*flaA* lux *L. pneumophila* from infected *Ifngr*^−/−^ and *Nos2*^−/−^ BMMs either not stimulated or stimulated with 6.0 ng/mL IFNγ. (**C**) RLU from LP02 Δ*flaA* lux *L. pneumophila* from infected *Myd88*^−/−^ and *Myd88*^−/−^ *Nod1*^−/−^ *Nod2*^−/−^BMMs either not stimulated or stimulated with 6.0 ng/mL IFNγ. (**D**) RLU from LP02 Δ*flaA* lux *L. pneumophila* from infected LysMCre^+^ *Atg5*^fl/fl^ and LysMCre^+^ BMMs either not stimulated or stimulated with 6.0 ng/mL IFNγ. (**E**) RLU from LP02 Δ*flaA* lux *L. pneumophila* from infected *Gbp*^chr3−/−^, *Gbp*^chr3^ ^+/+^, and *Casp1/11*^−/−^ BMMs either unstimulated or stimulated with 6.0 ng/mL IFNγ. Data reflect individual experiments that represent at least two independent experiments. Error bars in all graphs represent standard deviation of the mean of at least three technical replicates. p < 0.001 comparing no stim vs. IFNγ curves in all genotypes of BMMs (except *Ifngr*^−/−^) by 2-way ANOVA. No stim and IFNγ curves do not differ significantly in *Ifngr*^−/−^ BMMs.

### 2-deoxyglucose partially reverses IFNγ-dependent restriction of *L. pneumophila* in BMMs

We next investigated the possibility that IFNγ may act to restrict *L. pneumophila* not through induction of any single antimicrobial factor but by changing the metabolic landscape of the host macrophage to be unsuitable for the metabolic needs of *L. pneumophila*. As macrophages infected with *L. pneumophila* and IFNγ-stimulated macrophages increase rates of glycolysis (1, 36, 42), we tested whether treatment with the glycolysis inhibitor 2-deoxyglucose (2DG) might interfere with IFNγ-dependent restriction observed in BMMs. 2DG is metabolized to 2DG-phosphate in cells, which is directly antimicrobial (36). However, by taking advantage of a newly identified strain of *L. pneumophila* resistant to the direct antimicrobial effect of 2DG(P) in BMMs (Δ*flaA*Δ*uhpC L. pneumophila*) (36), we observed that addition of 2DG to BMMs partially restored *L. pneumophila* replication in IFNγ-treated macrophages (**Figure 2A**). We confirmed that 2DG disrupts the enhanced glycolysis observed in *L. pneumophila*-infected BMMs stimulated with IFNγ (**Figure 2B**).

**Figure 2.**
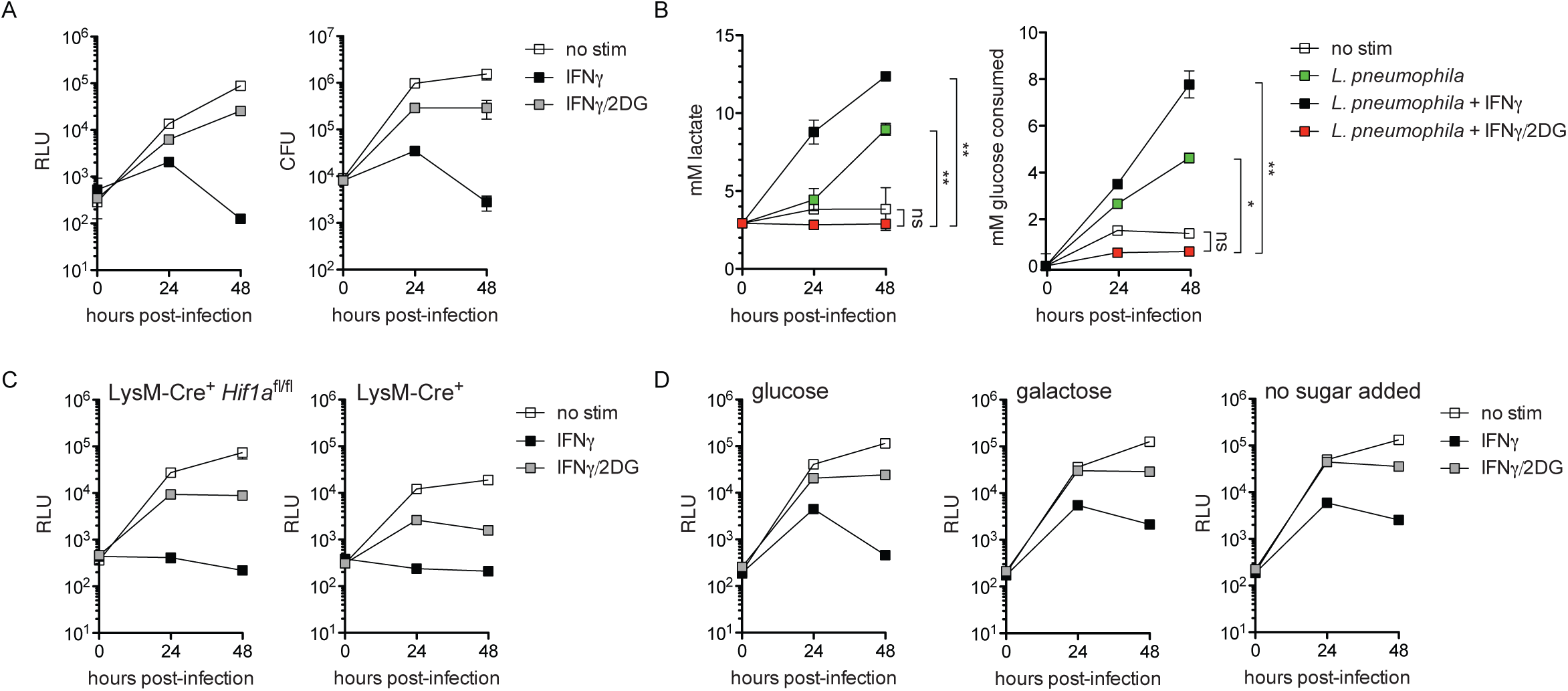
2DG rescues *L. pneumophila* replication in IFNγ-stimulated macrophages. (**A**) RLU (left) and CFU (right) of LP02 Δ*flaA*Δ*uhpC* lux *L. pneumophila* from infected WT C57BL/6 BMMs either not stimulated (no stim), stimulated with 6.0 ng/mL IFNγ (IFNγ), or stimulated with 6.0 ng/mL IFNγ + 2.0 mM 2DG (IFNγ/2DG). p < 0.001 comparing all curves to each other in each graph by 2-way ANOVA. (**B**) Lactate secretion (left) and glucose consumption (right) measured in cell culture media following infection of WT C57BL/6 BMMs with LP02 Δ*flaA*Δ*uhpC* lux *L. pneumophila* and stimulated with 6.0 ng/mL IFNγ and 1.0 mM 2DG as indicated. ** = p < 0.01; * = p < 0.05; ns = not significant comparing indicated curves by 2-way ANOVA. (**C**) RLU from LP02 Δ*flaA*Δ*uhpC* lux *L. pneumophila* from infected LysMCre^+^ *Hif1a*^fl/fl^ and LysMCre^+^ BMMs not stimulated or stimulated with 6.0 ng/mL IFNγ and 2.0 mM 2DG as indicated. p < 0.01 comparing all curves to each other corresponding to each genotype by 2-way ANOVA. (**D**) RLU from LP02 Δ*flaA*Δ*uhpC* lux *L. pneumophila* from infected WT C57BL/6 BMMs stimulated with 6.0 ng/mL IFNγ and 2.0 mM 2DG as indicated and cultured in infection media containing 11.11 mM glucose (left), 11.11 mM galactose in the absence of glucose (center), and in glucose-free media with no exogenous source of sugar (right). p < 0.001 comparing all curves to each other in each graph by 2-way ANOVA. Data reflect results of individual experiments that represent at least three independent experiments. Error bars in all graphs represent standard deviation of the mean of at least three technical replicates.

Previous studies have demonstrated that while *L. pneumophila* has the capacity to metabolize glucose, it does not rely on glucose or glucose derivatives to fuel its replication in broth, and is largely indifferent to perturbations in BMM glycolysis during infection (36, 43–45). To test whether induction of aerobic glycolysis by IFNγ restricts *L. pneumophila* replication, we performed infections using BMMs lacking hypoxia-inducible factor 1α (HIF1α), which fail to upregulate glycolysis in response to inflammatory stimuli and have a defect in IFNγ-mediated control of *Mtb* (14). We observed that HIF1α-deficient BMMs resembled wild-type BMMs in terms of IFNγ-dependent restriction and 2DG rescue of *L. pneumophila* replication (**Figure 2C**). Replacement of glucose with galactose, which inhibits increased glycolysis in IFNγ-stimulated BMMs (14, 46), also did not alter the ability of IFNγ to restrict or 2DG to rescue *L. pneumophila* replication (**Figure 2D**). Further, IFNγ was able to mediate bacteria restriction, and 2DG was able to reverse this restriction, in BMMs cultured in glucose-free media lacking any added sugar (**Figure 2D**). Finally, we tested whether other inhibitors of glycolysis, 3-bromopyruvate (3BP) and sodium oxamate (NaO), recapitulated the effects of 2DG. Neither 3BP nor NaO reversed IFNγ-dependent restriction of Δ*flaA*Δ*uhpC L. pneumophila* (the 2DG-resistant strain) or Δ*flaA L. pneumophila* (**Supplementary Figure 1**). Together, these data indicate that glycolysis inhibition is not required for IFNγ-mediated restriction of *L. pneumophila* replication in BMMs, and suggests that effects of 2DG other than glycolysis inhibition are responsible for its interference with the cell-intrinsic IFNγ-dependent immune response to *L. pneumophila* in BMMs.

### Some, but not all, unfolded protein response stimuli reverse IFNγ-dependent inhibition of *L. pneumophila*

To determine potential “off-target” effects of 2DG that could be responsible for reversal of IFNγ-mediated restriction of *L. pneumophila*, we performed transcript profiling on BMMs stimulated with the TLR2 agonist Pam3CSK4 or infected with *L. pneumophila*, stimulated with IFNγ ± 2DG. Pathway analysis of transcripts upregulated in 2DG conditions indicated induction of endoplasmic reticulum stress, also known as the unfolded protein response (UPR, **Supplementary Figure 2** and **Supplementary Table 1**). 2DG is thought to trigger the UPR due to interference with protein glycosylation pathways in the endoplasmic reticulum (47). This led us to hypothesize that induction of the UPR perturbs IFNγ-dependent restriction of *L. pneumophila* replication in BMMs. In fact, we observed that other drugs that trigger UPR stress, including geldanamycin, brefeldin A, and dithiothreitol also partially rescued *L. pneumophila* replication in IFNγ-stimulated BMMs (**Figure 3A**). However, not all drugs that trigger the UPR rescued *L. pneumophila* replication in IFNγ-treated BMMs. For example, treatment of BMMs with the potent UPR inducers tunicamycin or thapsigargin did not affect IFNγ-mediated restriction of *L. pneumophila* replication (**Figure 3B**).

**Figure 3.**
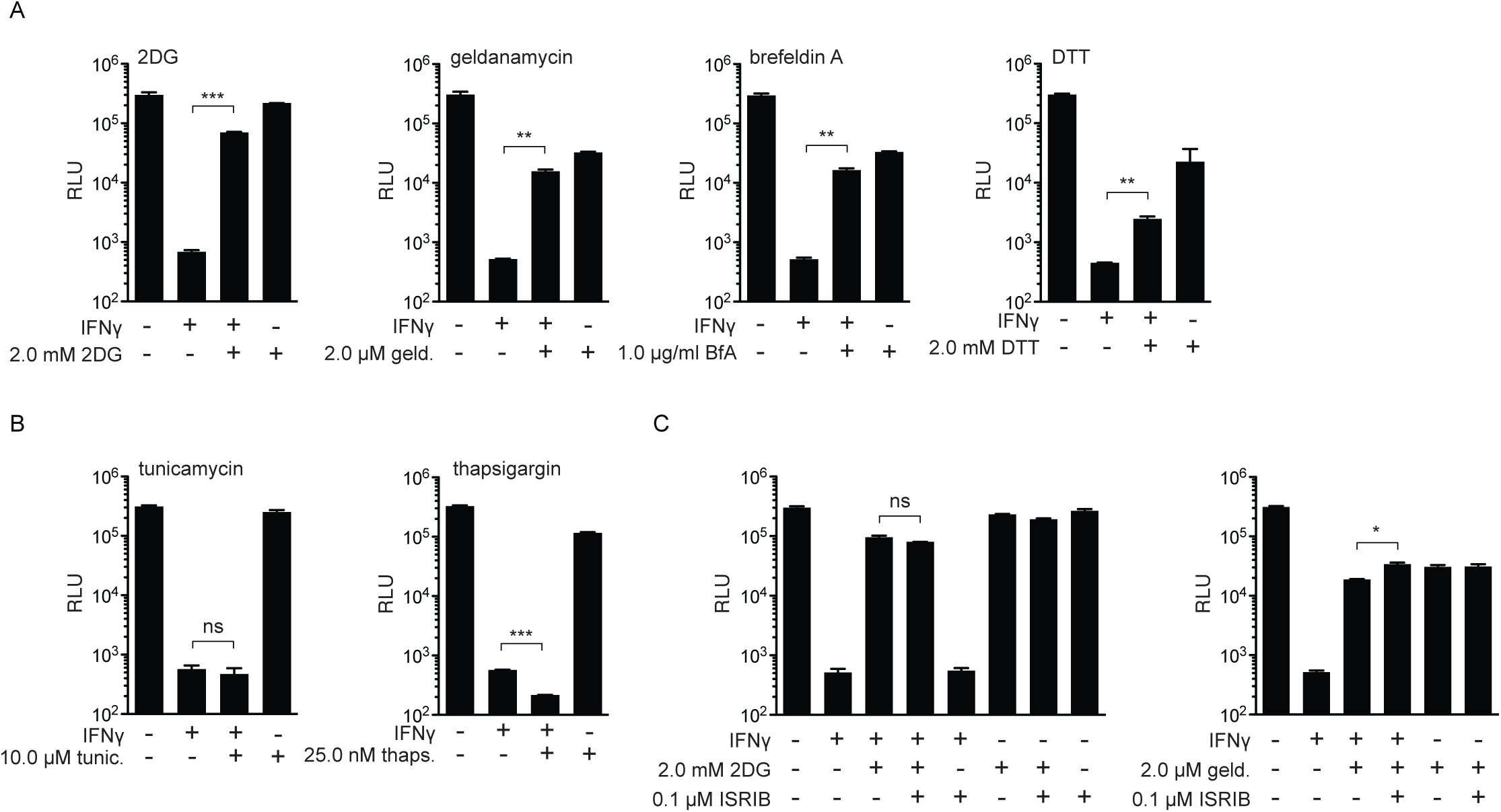
Differential effect of UPR stress stimuli on rescue of *L. pneumophila* replication in IFNγ-stimulated macrophages. (**A**) RLU from LP02 Δ*flaA*Δ*uhpC* lux *L. pneumophila* from infected WT C57BL/6 BMMs stimulated for 48 hours post-infection with 6.0 ng/mL IFNγ, 2.0 mM 2DG, 2.0 μM geldanamycin (geld.), 1.0 μg/ml brefeldin A (BfA), and 2.0 mM dithiothreitol (DTT) as indicated. (**B**) RLU from LP02 Δ*flaA*Δ*uhpC* lux *L. pneumophila* from infected WT C57BL/6 BMMs stimulated for 48 hours post-infection with 6.0 ng/mL IFNγ, 10.0 μM tunicamycin (tunic.), and 25.0 nM thapsigargin (thaps.) as indicated. (**C**) RLU from LP02 Δ*flaA*Δ*uhpC* lux *L. pneumophila* from infected WT C57BL/6 BMMs stimulated for 48 hours post-infection with 6.0 ng/mL IFNγ, 2.0 mM 2DG, 2.0 μM geldanamycin, and 0.1 μM ISRIB as indicated. *** = p < 0.001; ** = p < 0.01; * = p < 0.05; ns = not significant comparing means by unpaired *t* test. Data reflect results of individual experiments that represent at least three independent experiments. Error bars in all graphs represent standard deviation of the mean of at least two technical replicates. Concentrations of UPR stimuli displayed represent a single point in a titration at which we observed maximum effect on *L. pneumophila* replication in combination with IFNγ stimulation relative to a minimum effect on *L. pneumophila* replication in the absence of IFNγ.

One effect of UPR stress is arrest of protein translation via the PERK/EIF2α pathway, which can be reversed by the drug ISRIB (48). Importantly, ISRIB treatment did not interfere with 2DG- or geldanamycin-mediated rescue of *L. pneumophila* replication in IFNγ-stimulated BMMs, indicating that reversal of IFNγ-mediated restriction does not result from a global block in translation (**Figure 3C**). We confirmed that UPR stimuli and ISRIB were inducing UPR-associated transcripts and inhibiting ATF4-associated transcripts, respectively, via global transcript profiling (**Supplementary Figure 3B**) (49). Taken together, these results suggest that while some UPR-triggering drugs can partially reverse IFNγ-dependent restriction of *L. pneumophila* replication in BMMs, induction of the UPR does not inherently interfere with IFNγ-mediated restriction. Additionally, the partial rescue of bacterial replication in IFNγ-stimulated BMMs by UPR-triggering drugs does not act exclusively through general inhibition of protein translation.

### IFNγ fully restricts *L. pneumophila* in BMMs lacking IRG1, but is only partially restrictive in BMMs lacking both IRG1 and iNOS

Our analysis above revealed that certain UPR-stimulating drugs rescue *L. pneumophila* replication in IFNγ-stimulated BMMs, while others do not. We speculated that we could use these stimuli as a filter to look for transcripts associated with a restrictive vs. permissive macrophage state. Using this logic to filter results from RNAseq analysis of BMMs stimulated with Pam3CSK4 ± IFNγ ± UPR stimuli, we identified two genes, *Nos2* (encoding iNOS) and *Acod1* (encoding IRG1), whose transcript levels were elevated in restrictive conditions and lowered in permissive conditions (**Supplementary Figure 4A** and data not shown). Since iNOS deficiency has no effect on IFNγ-mediated control of *L. pneumophila* replication (e.g. Figure 1B), we speculated that IRG1 may restrict *L. pneumophila* replication in IFNγ-stimulated BMMs, as suggested (but not directly tested) previously (21). Using immortalized BMMs derived from C57BL/6 mice that inducibly express Cas9 (iCas9), we targeted *Acod1* and *Ifngr* with guide RNAs to generate BMMs that lack expression of IRG1 and IFNγ receptor, respectively (**Supplementary Figure 5** and **Table 1**). In comparison with *Ifngr*-targeted BMMs, which fail to restrict *L. pneumophila* when stimulated with IFNγ, we observed that *Acod1*-targeted immortalized BMMs fully retained the ability to restrict *L. pneumophila* replication upon stimulation with IFNγ (**Figure 4A**). We next generated primary BMMs from *Acod1*^−/−^ mice derived on the C57BL6/NJ background (26). Similar to immortalized BMMs, primary *Acod1*^−/−^ BMMs displayed intact IFNγ-dependent restriction of *L. pneumophila* (**Figure 4B**). These results suggest that IRG1 activity alone is not required for restriction of *L. pneumophila* in IFNγ-stimulated macrophages.

**Table 1.**
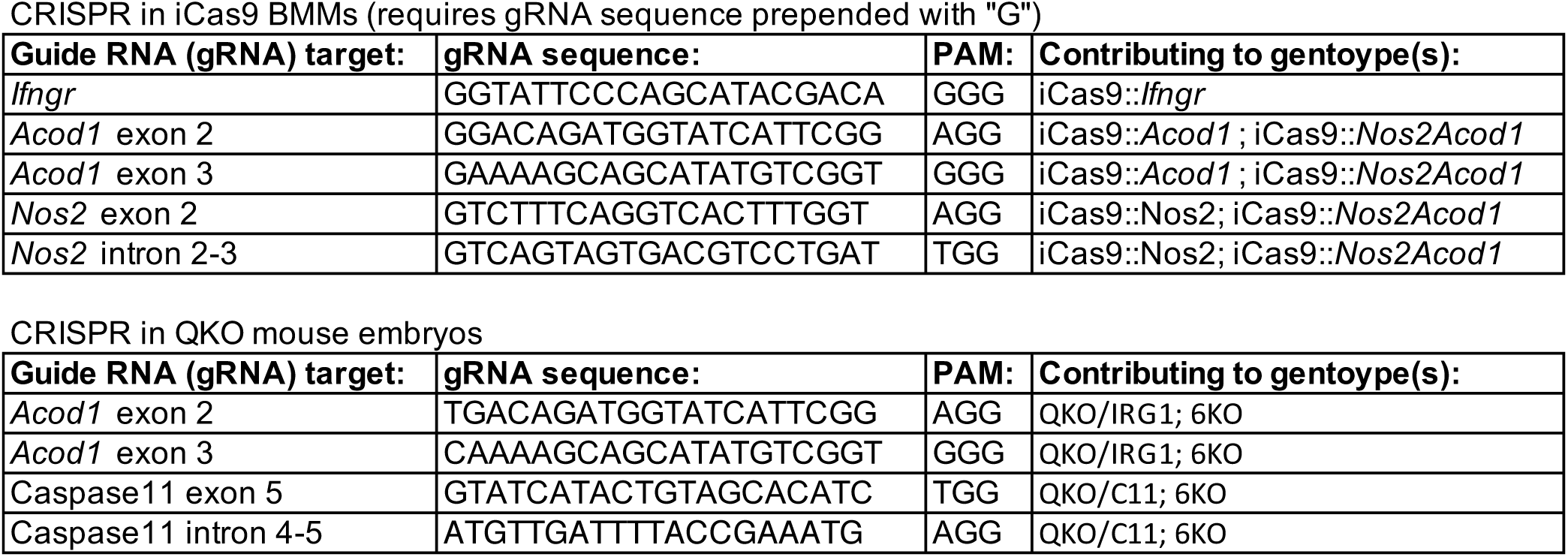
Guide RNAs used in this study.

**Figure 4.**
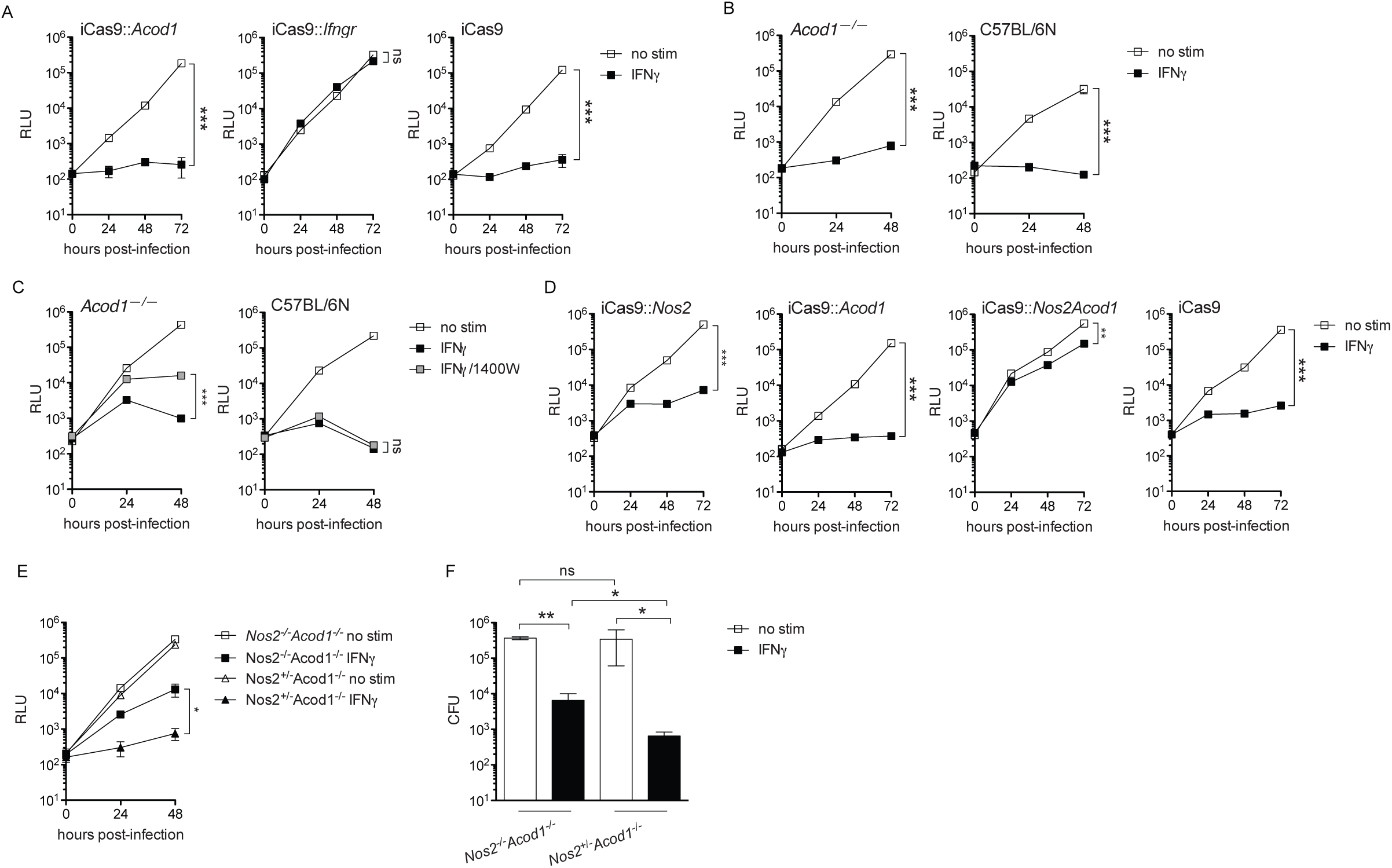
BMMs lacking IRG1 retain IFNγ-mediated restriction of *L. pneumophila* while BMMs lacking both INOS and IRG1 lose the ability to fully restrict *L. pneumophila*. (**A**) RLU from LP02 Δ*flaA* lux *L. pneumophila* from infected iCas9 BMMs in which *Acod1* was targeted with two guide RNAs (iCas9::*Acod1*), *Ifngr* was targeted with one guide RNA (iCas9::*Ifngr*), or that were not manipulated (iCas9) not stimulated (no stim) or stimulated with 6.0 ng/mL IFNγ. (**B**) RLU from LP02 Δ*flaA* lux *L. pneumophila* from infected primary *Acod1*^−/−^, and wild-type C57BL/6N BMMs not stimulated or stimulated with 6.0 ng/mL IFNγ. C57BL/6N BMMs were included as a wild-type control for BMMs derived from *Acod1*^−/−^ mice, which were generated on the C57BL/6N background. (**C**) RLU from LP02 Δ*flaA* lux *L. pneumophila* from infected *Acod1*^−/−^ and WT (C57BL/6N) BMMs either not stimulated (no stim), stimulated with 6.0 ng/mL IFNγ, or stimulated with 6.0 ng/mL IFNγ + 100 μM 1400W (IFNγ/1400W). p < 0.001 comparing no stim vs. IFNγ curves in both genotypes by 2-way ANOVA. IFNγ does not differ significantly from IFNγ+1400W in C57BL/6N BMMs. (**D**) RLU from LP02 Δ*flaA* lux *L. pneumophila* from infected iCas9 BMMs in which *Nos2* was targeted with two guide RNAs (iCas9::*Nos2*), *Acod1* was targeted with two guide RNAs (iCas9::*Acod1*), both *Nos2* and *Acod1* were targeted with two guide RNAs each (iCas9::*Nos2Acod1*), or that were not manipulated (iCas9) either not stimulated (no stim) or stimulated with 6.0 ng/mL IFNγ. (**E**) RLU from LP02 Δ*flaA* lux *L. pneumophila* from infected primary BMMs derived from *Nos2*^−/−^*Acod1*^−/−^ and littermate *Nos2*^+/−^ *Acod1*^−/−^ mice either not stimulated (no stim) or stimulated with 6.0 ng/mL IFNγ. *A* – *E*: *** = p < 0.001; ** = p < 0.01; * = p < 0.05; ns = not significant comparing indicated curves by 2-way ANOVA. (**F**) LP02 Δ*flaA* lux *L. pneumophila* CFU enumerated 48 hours post-infection in BMMs derived from *Nos2*^−/−^*Acod1*^−/−^ and littermate *Nos2*^+/−^*Acod1*^−/−^ mice either not stimulated (no stim) or stimulated with 6.0 ng/mL IFNγ. ** = p < 0.01; * = p < 0.05; ns = not significant comparing means by unpaired *t* test. Data reflect results of individual experiments that represent at least two independent experiments. Error bars in all graphs represent standard deviation of the mean of at least three technical replicates.

We next tested the hypothesis that iNOS and IRG1 activity are redundant in terms of effecting restriction of *L. pneumophila* in IFNγ-stimulated BMMs. In line with this hypothesis, we observed a partial (∼10-fold) loss of restriction in *Acod1*^−/−^ BMMs treated with the iNOS inhibitor 1400W, indicating that in the absence of IRG1, iNOS function is required to mediate full restriction of *L. pneumophila* in IFNγ-stimulated BMMs (**Figure 4C**). Reinforcing the idea that the activities of iNOS and IRG1 are redundant in terms of the IFNγ-coordinated response to *L. pneumophila*, we observed that targeting of both *Nos2* and *Acod1*, but not each factor independently, in iCas9 BMMs resulted in a marked loss of IFNγ-dependent restriction of *L. pneumophila* (**Figure 4D**). We next crossed *Nos2*^−/−^ and *Acod1*^−/−^ mice to derive littermate *Nos2*^−/−^ *Acod1*^−/−^ and *Nos2*^+/−^*Acod1*^−/−^ mice. We observed greater loss of IFNγ-dependent *L. pneumophila* restriction in *Nos2*^−/−^*Acod1*^−/−^ relative to *Nos2*^+/−^*Acod1*^−/−^ BMMs (**Figure 4E** and **4F**). In sum, these results indicate that the function of either iNOS or IRG1 must be intact to mediate full restriction of *L. pneumophila* replication in IFNγ-stimulated macrophages. This result suggests that activation of each of these factors by IFNγ is sufficient, individually, to mediate full restriction of *L. pneumophila*.

### BMMs deficient in six genes are fully defective in restriction *L. pneumophila* upon stimulation by IFNγ

While it appears that iNOS and IRG1 are sufficient and redundant in coordinating a large proportion of *L. pneumophila* restriction in IFNγ-stimulated BMMs, we observed that BMMs deficient in both iNOS and IRG1 retain partial restriction of *L. pneumophila* replication (**Figure 4D–4F**). In an effort to pinpoint the additional factors that mediate IFNγ-dependent restriction of *L. pneumophila* in BMMs, we made use of existing QKO mice lacking functional iNOS, NADPH oxidase 2 (NOX2), and immunity-related GTPase family M members 1 and 3 (IRGM1, IRGM3) (20). To test the hypothesis that the six factors implicated across our observations (iNOS, IRG1) and the studies by Pilla *et al* (QKO, CASP11) and Naujoks *et al* (IRG1) comprise the entirety of the IFNγ-coordinated response to *L. pneumophila* in macrophages, we employed CRISPR/Cas9 to target *Casp11* and *Acod1* (**Table 1**) in QKO mouse embryos to generate three novel mouse strains: QKO mice that also lack functional CASP11 (QKO/C11), QKO mice that also lack functional IRG1 (QKO/IRG1), and QKO mice that additionally lack both CASP11 and IRG1 (6KO). As previously reported, and in line with our observations in *Nos2* single knockout BMMs, we observed that IFNγ-dependent restriction of *L. pneumophila* in QKO BMMs was largely intact, indicating that no gene disrupted in these cells is absolutely required for restriction of *L. pneumophila* (**Figure 5A** and **5B**). QKO/C11 BMMs did not lose IFNγ-mediated restriction relative to QKO BMMs (**Figure 5A** and **5B**). In contrast, we observed a striking loss of restriction in QKO/IRG1 BMMs and, effectively, a total loss of *L. pneumophila* restriction in 6KO BMMs stimulated with IFNγ (**Figure 5A** and **5B**).

**Figure 5.**
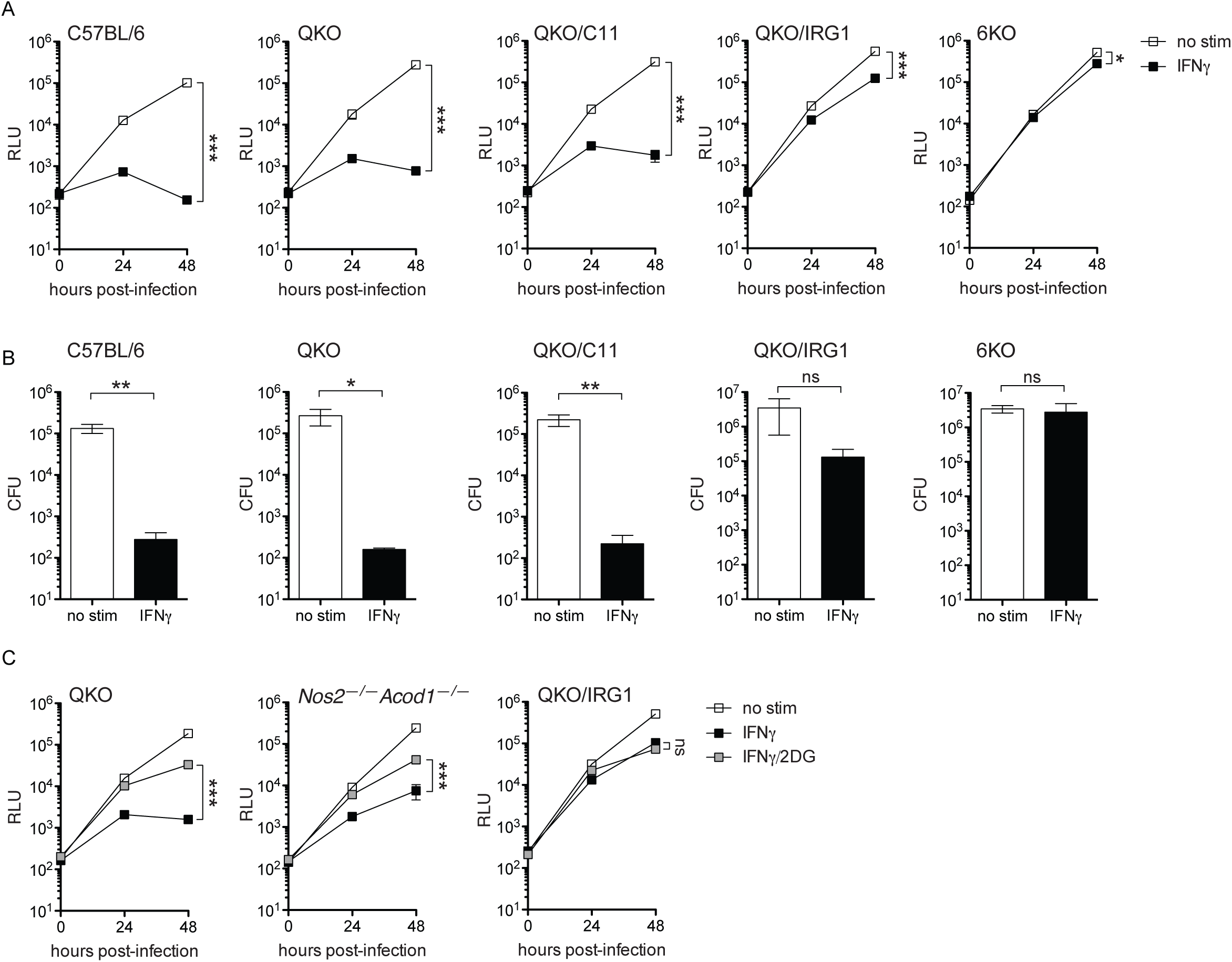
QKO BMMs that additionally lack functional either CASP11 and IRG1 or both factors display partial to full lack of restriction of *L. pneumophila* replication when stimulated with IFNγ. (**A**) RLU from LP02 Δ*flaA* lux *L. pneumophila* from infected C57BL/6, QKO, QKO/C11, QKO/IRG1, and 6KO BMMs either not stimulated (no stim) or stimulated with 6.0 ng/mL IFNγ. *** = p < 0.001; * = p < 0.05 comparing indicated curves by 2-way ANOVA. (**B**) CFU recovered from WT, QKO, QKO/C11, QKO/IRG1, and 6KO BMMs 48 hours following infection with LP02 Δ*flaA* lux *L. pneumophila*. **; = p < 0.01; * = p < 0.05; ns = not significant comparing means by unpaired *t* test. (**C**) RLU from LP02 Δ*flaA*Δ*uhpC* lux *L. pneumophila* from infected QKO, *Nos2*^−/−^*Acod1*^−/−^, and QKO/IRG1 BMMs stimulated with 6.0 ng/mL IFNγ and 2.0 mM 2DG as indicated. *** = p < 0.001**; ns = not significant comparing indicated curves by 2-way ANOVA. Data reflect results of individual experiments that represent at least three independent experiments. Error bars in all graphs represent standard deviation of the mean of at least three technical replicates.

If the ability of 2DG to rescue *L. pneumophila* in IFNγ-treated BMMs acts through inhibition of iNOS and IRG1, we would expect 2DG to have no effect in BMMs lacking expression of these factors. In fact, we observed that in comparison with QKO BMMs, 2DG retained the ability to partially rescue *L. pneumophila* replication in *Nos2*^−/−^ *Acod1*^−/−^ BMMs stimulated with IFNγ (**Figure 5C**). However, the rescue effect of 2DG was absent in QKO/IRG1 BMMs (**Figure 5C**). Transcript profiling did not reveal an inhibitory effect of 2DG on expression of the other genes disrupted in the QKO/IRG1 background (**Supplementary Figure 4B**). This result indicates that 2DG may mediate some beneficial metabolic effect for *L. pneumophila* independent of regulating activity of iNOS and IRG1; however, these effects may require the other factors disrupted in the QKO background. Alternately, the potential metabolic effects of 2DG may be obscured by the profound loss of restriction observed in QKO/IRG1 BMMs.

In sum, these results further underscore our previous observation that iNOS and IRG1 are each sufficient in terms of mediating a large proportion of the IFNγ-dependent restriction of *L. pneumophila* in BMMs, as the addition of IRG1 deficiency to the QKO background profoundly disabled IFNγ-mediated restriction. Further, our data reveal a partial role for CASP11 in control of *L. pneumophila* restriction in IFNγ-stimulated BMMs, given the differences observed between QKO/IRG1 and 6KO BMMs. Finally, our results demonstrate that the six genes disrupted in 6KO BMMs, or a subset of those six that includes *Nos2*, *Acod1*, and *Casp11*, coordinate(s) the entirety of the IFNγ-dependent, cell-intrinsic control of *L. pneumophila* observed in BMMs.

## Discussion

Our results support a model in which IFNγ restricts *L. pneumophila* replication in mammalian macrophages through activation of multiple redundant factors including iNOS and IRG1. To date, no study has identified a macrophage deficient in any single IFNγ-stimulated gene that is impaired in the ability to restrict *L. pneumophila* replication when stimulated with IFNγ. Even QKO macrophages, which lack three other potentially antimicrobial factors in addition to iNOS, do not lose IFNγ-mediated restriction of *L. pneumophila*, reinforcing the notion that redundant mechanisms contribute to IFNγ-mediated bacterial control in macrophages.

Our recent identification of a strain of *L. pneumophila* resistant to the direct antimicrobial effect of 2DG when growing in BMMs (36) allowed us to test the hypothesis that global disruption of macrophage metabolism would interfere with the antimicrobial effects of IFNγ. Indeed, 2DG partially reversed the restriction of *L. pneumophila* replication by IFNγ in BMMs. However, neither the glycolysis inhibition activity nor the UPR induction activity of 2DG, *per se*, appears to underlie the ability of this drug to subvert the antimicrobial effect of IFNγ. Instead, 2DG appears to regulate the IFNγ-dependent induction of iNOS and IRG1 via some as-yet unidentified mechanism. In addition, there appears to be some effect of 2DG independent of iNOS and IRG1 regulation, suggesting that metabolic perturbation could interfere with the antimicrobial activities of IFNγ in infected macrophages. Ultimately, experimentation with 2DG and other stimuli that reversed IFNγ-mediated restriction of *L. pneumophila* led us to the discovery that both iNOS and IRG1 appear to be fully sufficient, and therefore redundant, in terms of mediating IFNγ-coordinated immune response to *L. pneumophila* in macrophages.

A complex picture is emerging in terms of the role of IRG1 and the metabolite it produces, itaconate, during inflammation and infection. A direct antimicrobial role for itaconate via poisoning the bacterial glyoxylate pathway has been suggested for *Mtb* and *L. pneumophila* (21, 25). IRG1 was shown to be an essential component of the immune response to *Mtb*, as *Acod1*^−/−^ mice succumbed more rapidly than wild-type to infection; however, IRG1 appeared to be required for regulation of non-cell-autonomous pathological inflammation, and there was no evidence for cell-intrinsic antimicrobial effects of itaconate (50). IRG1 has also been demonstrated to be protective in a model of Zika virus infection in neurons (51). Interestingly, other studies have demonstrated anti-inflammatory effects of itaconate on myeloid cells, suggesting it may act as part of a negative feedback loop to control inflammation (27, 52). Beyond production of itaconate, the disruption of oxidative metabolic pathways caused by IRG1 activity may promote antimicrobial metabolic shifts in macrophages. Ultimately, diverse cell-intrinsic and intercellular roles for IRG1 and itaconate likely contribute to the immune response to a broad array of pathogens. Our data demonstrate that IRG1 is not required for the cell-intrinsic immune response to *L. pneumophila* in macrophages treated with IFNγ, but indeed may be sufficient to coordinate IFNγ-mediated restriction of *L. pneumophila,* as previously reported (21). Both NO generated by iNOS and itaconate generated by IRG1 may be directly antimicrobial to *L. pneumophila* in macrophages stimulated with IFNγ. Alternately or additionally, iNOS and IRG1 may act to restrict *L. pneumophila* replication via coordinating global changes in macrophage metabolism that restrict access to key bacterial metabolites or otherwise render the host macrophage inhospitable for bacterial growth.

Adding *Acod1* and *Casp11* deficiency to the QKO background revealed further layers of redundancy in the immune response to *L. pneumophila* coordinated by IFNγ. While QKO/C11 macrophages did not differ meaningfully from QKO macrophages in terms of IFNγ-mediated bacterial restriction, we observed a profound loss of restriction in QKO/IRG1 macrophages, beyond what we observed in primary *Nos2*^−/−^ *Acod1*^−/−^ macrophages. This result indicates that factors other than iNOS disrupted in the QKO background may play a role in limiting *L. pneumophila* in IFNγ-stimulated macrophages.

In agreement with the results of Pilla *et al* (20), our data suggest that a role exists for CASP11 in the IFNγ-mediated immune response to *L. pneumophila*, given the near-complete inability of IFNγ to restrict *L. pneumophila* replication in 6KO macrophages vs. QKO/IRG macrophages (that retain CASP11). In combination with the data showing that *Casp1/11*^−/−^ and QKO/C11 BMMs retain IFNγ-mediated bacterial restriction, this result demonstrates that the activity of CASP11 is also redundant, at least with the activities of iNOS and IRG1.

In sum, our study reveals a more comprehensive picture of the factors that are necessary and sufficient to coordinate the IFNγ-dependent immune response to *L. pneumophila*. While we have not determined whether all six of the genes disrupted in 6KO BMMs cells are required to fully exert IFNγ-dependent cell-intrinsic restriction of *L. pneumophila* or a subset of the six that includes iNOS, IRG1, and CASP11, we are encouraged that among the numerous genes transcribed in IFNγ-stimulated macrophages we have narrowed the field that mediate cell-intrinsic control of *L. pneumophila* to six candidates. While all of the gene products disrupted in the 6KO background could function directly as antimicrobial effectors, we also note the possibility that some or all may function as upstream regulators and thus affect *L. pneumophila* indirectly.

IFNγ is an essential component of the immune response to bacterial pathogens beyond *L. pneumophila*. Thus, the implications of this study extend beyond furthering our understanding of the immune response to *L. pneumophila*, an accidental pathogen of mammals that did not evolve to evade the human immune response. Our work reveals fundamental redundancy in the IFNγ-dependent immune response to potentially pathogenic environmental microbes. Dissecting these overlapping innate immune strategies reveals the complexity and comprehensiveness of the innate immune barrier posed to novel environmental microorganisms by mammalian macrophages and IFNγ. Further, a more detailed understanding of how IFNγ can mediate bacterial restriction in host cells may inform studies of how “professional” pathogens, such as *Mtb*, *S. enterica*, and *L. monocytogenes*, have evolved to avoid or subvert these effects of IFNγ.

## Materials and Methods

### Ethics statement

We conducted experiments in this study according to guidelines established by the *Guide for the Care and Use of Laboratory Animals* of the National Institutes of Health (53) under a protocol approved by the Animal Care and Use Committee at the University of California, Berkeley (AUP-2014-09-6665).

### Bone marrow-derived macrophages

We purchased wild-type C57BL/6 (strain 000664), *Ifngr*^−/−^ (strain 003288), *Ifnar*^−/−^ (strain 028288), *Myd88*^−/−^ (strain 009088), *Nos2*^−/−^ (strain 002609), *Acod1*^−/−^ (strain 029340), and C57BL/6N (strain 005304) mice from Jackson Laboratory as a source of bone marrow to derive macrophages. *Casp1/11*^−/−^ mice were provided by A. Van der Velden and M. Starnbach (54). *Myd88*^−/−^ *Nod1*^−/−^ *Nod2*^−/−^ mice were generated at UC Berkeley as described previously (39). *Nos2*^−/−^*Irg1*^−/−^, and *Nos2*^+/−^*Irg1*^−/−^ mice were also generated by crossing in-house at UC Berkeley. QKO mice (*Nos2*^−/−^, *Cybb*^−/−^, *Irgm1*^−/−^, *Irgm3*^−/−^), generously provided by the lab of Christopher Sassetti at the University of Massachusetts, and mice lacking a section of chromosome 3 containing GBPs 1, 2, 3, 5, and 7 (*Gbp*^chr3–/−^) and wild-type control mice (*Gbp*^chr3+/+^), all on the C57BL/6 background were generated as described (20). We derived bone marrow-derived macrophages (BMMs) in RPMI supplemented with 10% fetal bovine serum, 2.0 mM L-glutamine, 100 μM streptomycin (all from Life Technologies), and 5% supernatant from 3T3 cells expressing macrophage colony-stimulating factor (generated in-house). Macrophages derived from LysMCre^+/+^ and LysMCre^+/+^*Atg5*^fl/fl^ mice on the C57BL/6 background were provided by Daniel Portnoy and Jeffery Cox at UC Berkeley. Macrophages derived from LysMCre^+/+^ and LysMCre^+/+^*Hif1a*^fl/fl^ mice on the C57BL/6 background were provided by Sarah Stanley at UC Berkeley.

### Mouse CRISPR

We generated QKO/C11 and QKO/IRG1 mice by pronuclear injection of Cas9 mRNA and guide RNAs into fertilized embryos of QKO mice as described previously (55). Founder male mice heterozygous for mutation in either *Casp11* or *Acod1* were backcrossed once onto the QKO background and offspring were intercrossed to generate QKO/C11, QKO/IRG1, and 6KO mice. *Acod1* mutation was determined by amplifying a fragment of genomic DNA surrounding the cut site targeted in *Acod1* exon 2 (forward primer: AACTCTGGGAATGCCAGCTC, reverse primer: GGAGCCACAACAGGGATCAA, yielding a ∼440 base-pair PCR product) and Sanger sequencing, which revealed a three-nucleotide deletion (TTC) and a one-nucleotide insertion (A) at the cut site in mutant DNA resulting in a frame-shift mutation and premature stop codon. *Casp11* mutation was determined by amplifying genomic DNA surrounding the cut sites indicated by both guide RNAs (forward primer: GGGGCTCTGAAAAGGTGTGA, reverse primer: TCTAGACACAAAGCCCATGT, revealing a ∼520-base pair band in wild-type DNA and a ∼290-base pair band in mutant DNA, indicating a missing ∼230-base pair fragment in mutant genomic DNA.

### iCas9 CRISPR

We cloned template DNA for the indicated guide RNAs into a pLX-sgRNA construct additionally containing blasticidin resistance (Addgene plasmid #50662). We transfected constructs into HEK293T cells along with lentivirus packaging vector pSPAX2 (Addgene plasmid #12260) and lentivirus envelope vector VSV-G (Addgene plasmid #8454). We used the resulting virus particles to transduce immortalized wild-type C57BL/6 cells that express doxycycline-inducible SpCas9 enzyme (generated using Addgene plasmid #50661). We cultured transduced cells in 3.0 μg/ml blasticidin (Invivogen) and 5.0 μg/ml doxycycline (Sigma) for at least two weeks prior to use in experiments.

### Bacterial strains, infection and stimulation of BMMs

LP02 is a thymidine auxotroph derived from LP01, a clinical isolate of *L. pneumophila* (56). Generation of Δ*flaA* and luminescent strains of *L. pneumophila* have been described previously (36, 37). We cultured all strains of *L. pneumophila* in AYE (ACES-buffered yeast extract broth) or on ACES-buffered charcoal-yeast extract (BCYE) agar plates at 37 °C. For measurement of intracellular *L. pneumophila* growth by luminescence or by CFU, we plated 100k BMMs/well in opaque white TC-treated 96-well microtiter plates and infected with *L. pneumophila* at a multiplicity of infection of 0.05. One hour post-infection by centrifugation at 287 × *g*, we replaced the media of infected BMMs with media ± stimulation at indicated concentrations. At the indicated times following infection, we measured bacterial growth by detection of luminescence at λ = 470 using a Spectramax L luminometer (Bio-Rad) or by dilution of infected cultures on BYCE agar plates for enumeration of CFU. Pam3CSK4 and *E. coli*-derived LPS were purchased from Invivogen. As indicated, we added recombinant mouse IFNγ (ThermoFisher), 2-deoxyglucose (Abcam), brefeldin A (BD), 1400W (Cayman Chemical), 3-bromopyruvte, sodium oxamate, galactose, geldanamycin, dithiothreitol, tunicamycin, thapsigargin, and ISRIB (all from Sigma). We performed lactate and glucose measurement with kits purchased from Sigma according to manufacturer instructions.

### Western Blot

Following stimulation for 24 hours, we mixed lysates derived from 1.0 × 10^6^ BMMs per stimulation condition with SDS sample buffer (40% glycerol, 8% SDS, 2% 2-mercaptoethanol, 40 mM EDTA, 0.05% bromophenol blue and 250 mM Tris-HCl, pH 6.8), boiled for 5 min and then separated by SDS-PAGE. Rabbit Anti-IRG1 antibody was from Abcam (ab222417) and mouse anti-β actin antibody was from Santa Cruz (47778).

### RNAseq

We submitted RNA purified from the indicated cell culture conditions using an RNeasy kit (Qiagen) to the QB3-Berkeley Functional Genomics Laboratory, where single-read 100 base pair read length (SR100) sequencing libraries were generated. Libraries were sequenced using either a HiSeq2500 System (Illumina) at the New York Genome Center (New York, NY) or a HiSeq4000 System (Illumina) at the Vincent J. Coates Genomics Sequencing Laboratory at UC Berkeley. We performed alignment, differential expression analysis, and gene set enrichment as described previously (57–59).

### Data Availability

We deposited the RNAseq data associated with this study in the NCBI Gene Expression Omnibus, available at https://www.ncbi.nlm.nih.gov/geo/ via accession numbers GSE135385 and GSE135386.

## Acknowledgements

R.E.V. is supported by an Investigator Award from the Howard Hughes Medical Institute and by NIH grants AI063302 and AI075039. We would like to thank Harmandeep Dhaliwal at the Cancer Research Laboratory Gene Targeting Facility at UC Berkeley for assistance in generating mouse strains used in this study. We would also like to thank Kevin Barry for assistance with RNAseq analysis. We acknowledge stimulating discussions with Sarah Stanley, Jonathan Braverman, Greg Barton, Daniel Portnoy, and members of the Vance, Stanley, Barton, and Portnoy Labs, and members of the P01 Intracellular Pathogens and Innate Immunity research group. The authors have no conflicts of interest with regard to the results presented in this study.

---

**Supplementary Figure 1.**
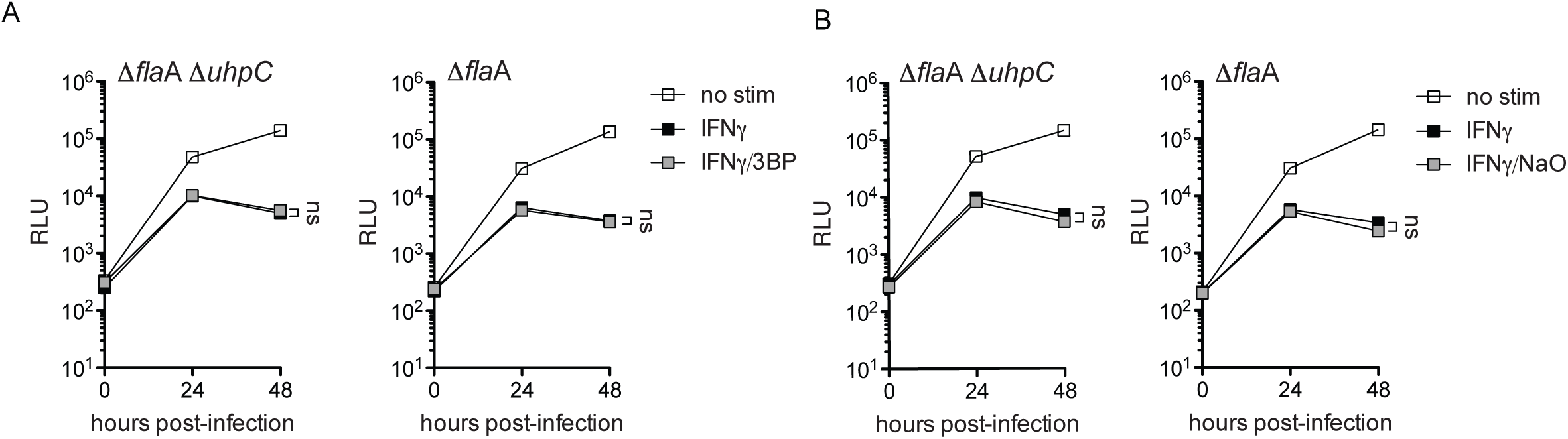
3-bromopyruvate and sodium oxamate do not rescue ΔflaAΔuhpC L. pneumophila replication in IFNγ-stimulated BMMs. (**A**) RLU from LP02 Δ*flaA*Δ*uhpC* lux *L. pneumophila* (left) and Δ*flaA* lux *L. pneumophila* (right) from infected WT C57BL/6 BMMs stimulated with 6.0 ng/mL IFNγ ± 60.0 μM 3-bromopyruvate (3BP). (**C**) RLU from LP02 Δ*flaA*Δ*uhpC* lux *L. pneumophila* (left) and LP02 Δ*flaA* lux *L. pneumophila* (right) from infected WT C57BL/6 BMMs stimulated with 6.0 ng/mL IFNγ and 2.5 mM sodium oxamate (NaO). Data reflect results of individual experiments that represent at least three independent experiments. Error bars in all graphs represent mean ± standard deviation of at least two technical replicates. Concentrations of 3BP and NaO displayed represent the highest single point in a titration at which we observed a minimum effect on *L. pneumophila* replication in the absence of IFNγ.

**Supplementary Figure 2.**
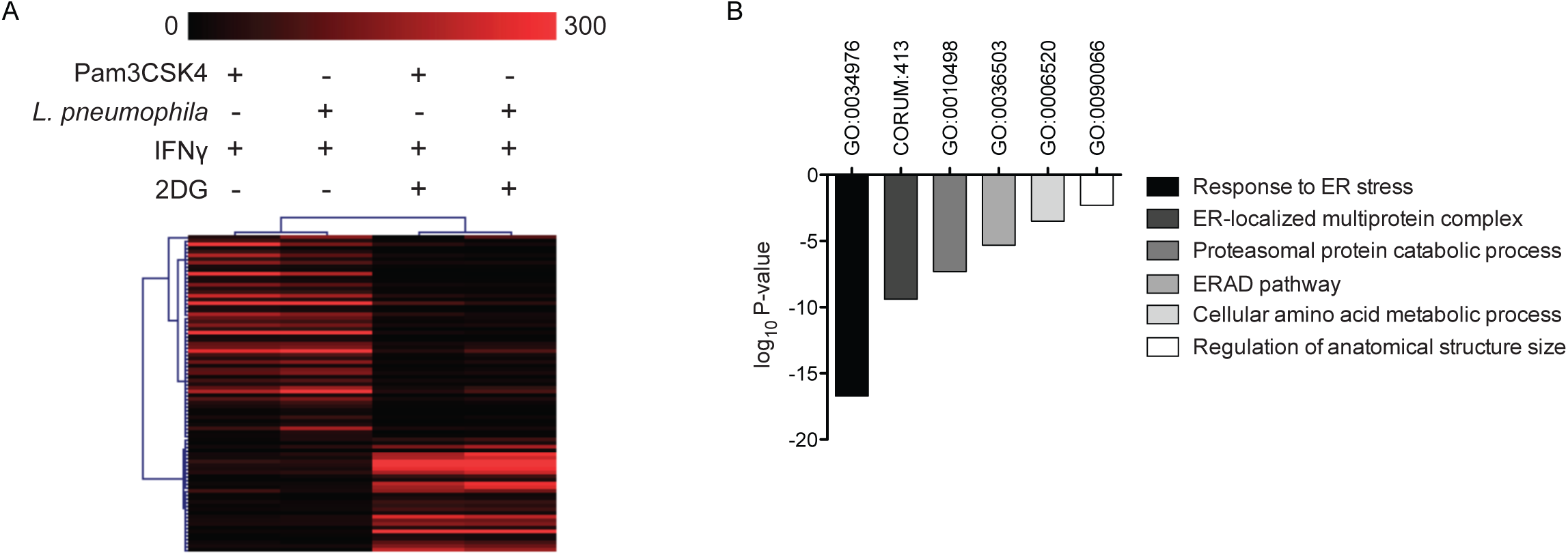
2DG triggers a transcriptional profile indicating endoplasmic reticulum stress in IFNγ-stimulated BMMs. (**A**) Heat map showing fragments per kilobase million (FPKM) of differentially expressed transcripts measured by RNAseq recovered from 1.0 × 10^6^ C57BL/6 BMMs/condition stimulated for 18 hours with 50.0 ng/ml Pam3CSK4 + 2.0 ng/ml IFNγ ± 1.5 mM 2DG and 1.0 × 10^6^ C57BL/6 BMMs/condition infected at T_0_ with LP02 Δ*flaA L. pneumophila* and stimulated 1 hour post-infection with 2.0 ng/ml IFNγ ± 1.5 mM 2DG for a total of 18 hours prior to harvest. Transcript IDs, FPKM, and log_2_ fold-change are shown in Supplementary Table 1. Only transcripts shown to differ significantly between both Pam3CSK4/IFNγ and *L.p*./IFNγ ± 2DG conditions, as calculated using TopHat/Cufflinks (57) are displayed. (**B**) Log_10_ P-value of gene ontology (GO) and comprehensive resource of mammalian protein complexes (CORUM) gene sets in which transcripts significantly upregulated in + 2DG conditions are enriched as determined using Metascape (59).

**Supplementary Figure 3.**
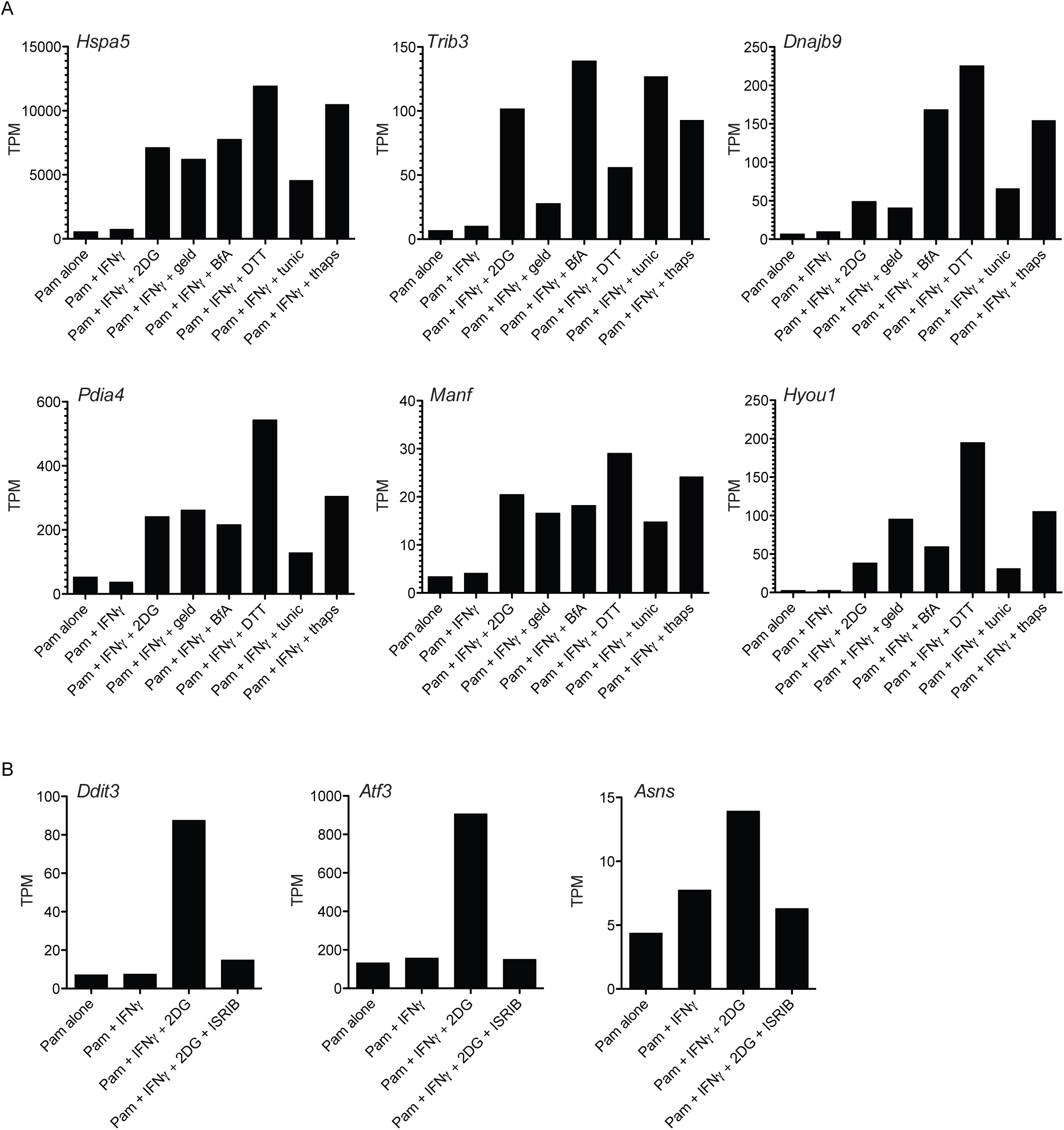
Validation of UPR gene expression by UPR stimuli and inhibition of ATF4-dependent gene expression by ISRIB. (**A**) Histograms displaying transcripts per million (TPM) of *Hspa5* (heat shock protein 5, aka BIP), *Trib3* (tribbles pseudokinase 3), *Dnajb3* (DnaJ heat shock protein family (Hsp40) member B3), *Pdia4* (protein disulfide isomerase associated 4), *Manf* (mesencephalic astrocyte-derived neurotrophic factor), and *Hyou1* (hypoxia up-regulated 1) measured by RNAseq recovered from 1.0 × 10^6^ BMMs/condition stimulated for 18 hours with 100 ng/ml Pam3CSK4 (Pam) alone or in combination with 6.0 ng/ml IFNγ, IFNγ + 2.0 mM 2DG, IFNγ + 2.0 μM geldanamycin (Geld), IFNγ + 1.0 μg/ml brefeldin A (BfA), IFNγ + 2.0 mM dithiothreitol (DTT), IFNγ + 10.0 μM tunicamycin (Tunic), or IFNγ + 25.0 nM thapsigargin (Thaps). These genes are a subset associated with gene ontology term GO:0034976, response to endoplasmic reticulum stress. (**B**) Histograms displaying TPM of *Ddit3* (DNA-damage inducible transcript 3, aka CHOP), *Atf3* (activating transcription factor 3), and *Asns* (asparagine synthetase) measured by RNAseq recovered from 1.0 × 10^6^ BMMs/condition stimulated for 18 hours with 100 ng/ml Pam3CSK4 (Pam) alone or in combination with 6.0 ng/ml IFNγ, IFNγ + 2.0 mM 2DG, or IFNγ + 2.0 mM 2DG + 0.1 μM ISRIB. These genes are associated with PERK/ATF4-dependent gene expression (49).

**Supplementary Figure 4.**
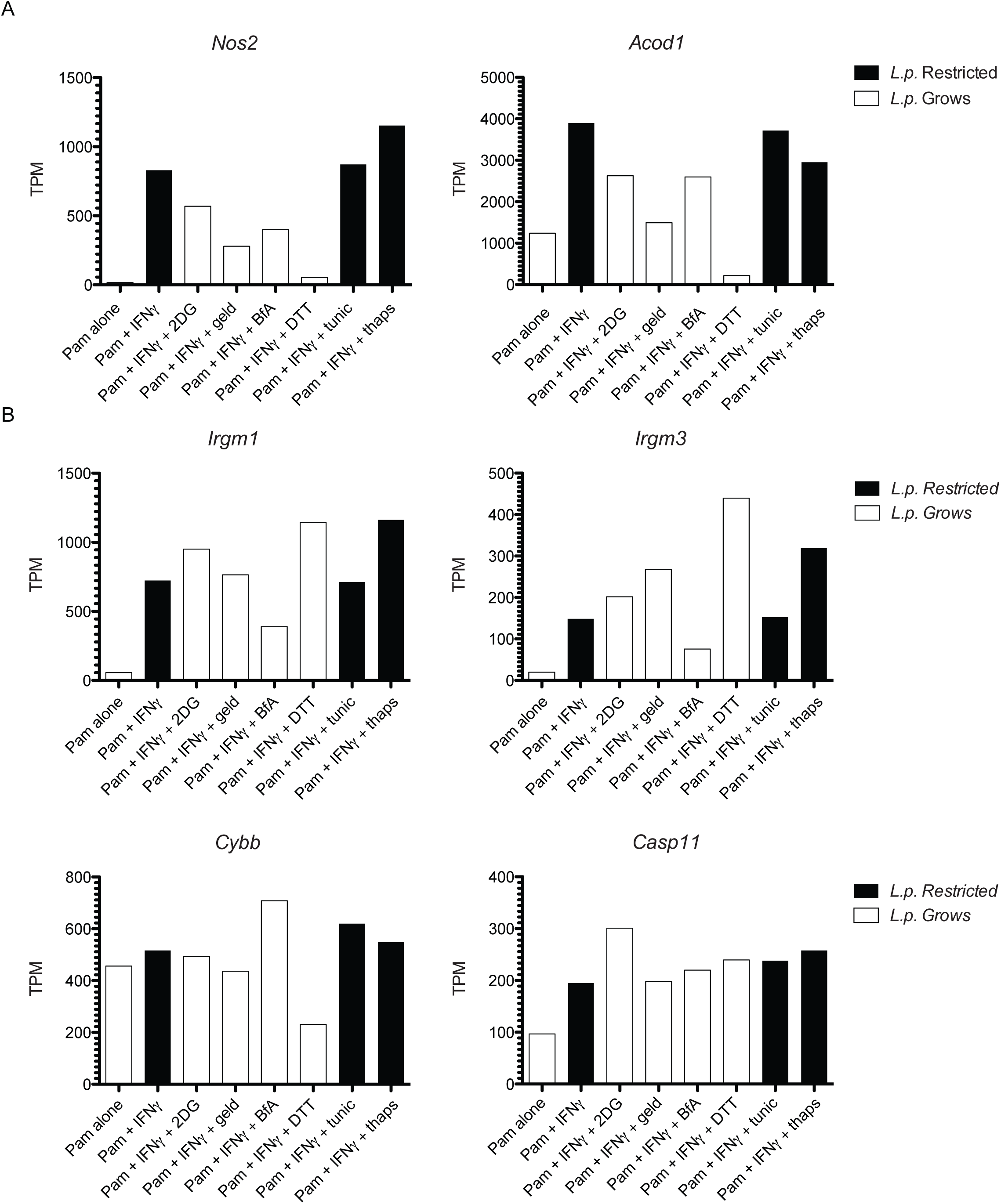
Nos2 and Acod1 transcription segregates with conditions permissive and restrictive for L. pneumophila replication in BMMs. (**A** and **B**) Histograms displaying transcripts per million (TPM) of *Nos2*, *Acod1*, *Irgm1*, *Irgm3*, *Cybb*, and *Casp11* measured by RNAseq recovered from 1.0 × 10^6^ BMMs/condition stimulated for 18 hours with 100 ng/ml Pam3CSK4 (Pam) alone or in combination with 6.0 ng/ml IFNγ, IFNγ + 2.0 mM 2DG, IFNγ + 2.0 μM geldanamycin (Geld), IFNγ + 1.0 μg/ml brefeldin A (BfA), IFNγ + 2.0 mM dithiothreitol (DTT), IFNγ + 10.0 μM tunicamycin (Tunic), IFNγ + 25.0 nM thapsigargin (Thaps).

**Supplementary Figure 5.**
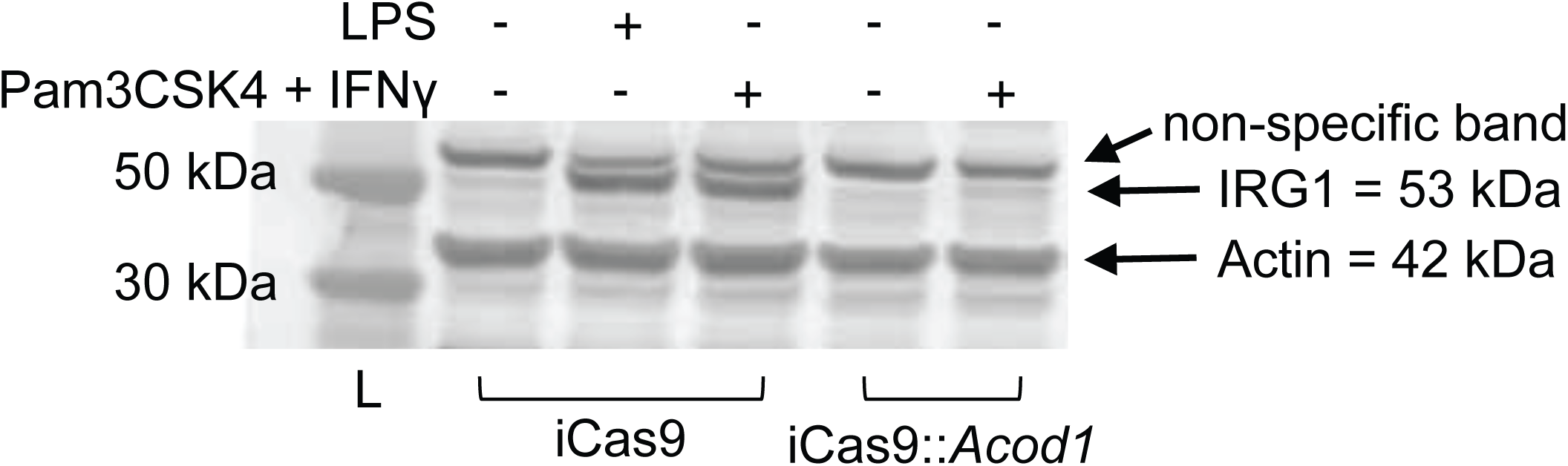
Validation of CRISPR-mediated targeting of Acod1 in iCas9 BMMs. Western blot demonstrating loss of IRG1 protein expression in iCas9::*Acod1* BMMs when stimulated for 24 hours with either 100 ng/ml *E. coli* lipopolysaccharide (LPS) or 100 ng/ml Pam3CSK4 + 10.0 ng/ml IFNγ. IRG1 migrates at ∼53 kDa; the IRG1 antibody also stains a non-specific band at a slightly higher molecular weight. A separate antibody was used to stain for mouse β actin, which migrates at ∼42 kDa.

**Supplementary Table 1.**
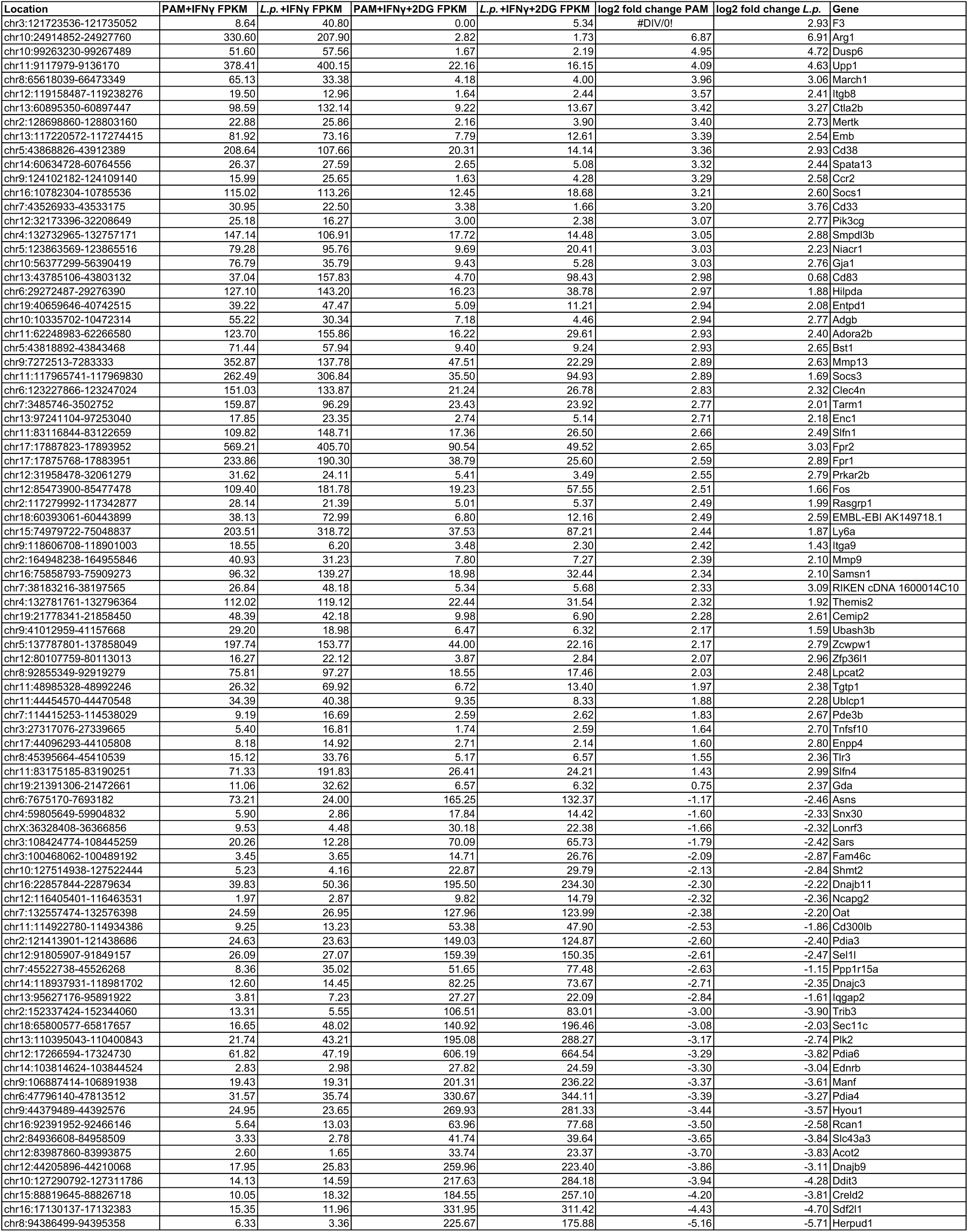
Transcripts that vary significantly in BMMs treated with Pam3CSK4 + IFNγ vs. Pam3CSK4 + IFNγ + 2DG.

